# Vocal markers of autism: assessing the generalizability of machine learning models

**DOI:** 10.1101/2021.11.22.469538

**Authors:** Astrid Rybner, Emil Trenckner Jessen, Marie Damsgaard Mortensen, Stine Nyhus Larsen, Ruth Grossman, Niels Bilenberg, Cathriona Cantio, Jens Richardt Møllegaard Jepsen, Ethan Weed, Arndis Simonsen, Riccardo Fusaroli

**Author notes:** **Corresponding author:** Riccardo Fusaroli, Aarhus University. N.B. indicates shared first authorship.

## Abstract

Machine learning (ML) approaches show increasing promise in their ability to identify vocal markers of autism. Nonetheless, it is unclear to what extent such markers generalize to new speech samples collected e.g., using a different speech task or in a different language. In this paper, we systematically assess the generalizability of ML findings across a variety of contexts.

We train promising published ML models of vocal markers of autism on novel cross-linguistic datasets following a rigorous pipeline to minimize overfitting, including cross-validated training and ensemble models. We test the generalizability of the models by testing them on i) different participants from the same study, performing the same task; ii) the same participants, performing a different (but similar) task; iii) a different study with participants speaking a different language, performing the same type of task.

While model performance is similar to previously published findings when trained and tested on data from the same study (out-of-sample performance), there is considerable variance between studies. Crucially, the models do not generalize well to different, though similar, tasks and not at all to new languages. The ML pipeline is openly shared.

Generalizability of ML models of vocal markers of autism is an issue. We outline three recommendations for strategies researchers could take to be more explicit about generalizability and improve it in future studies.

**LAY SUMMARY:** Machine learning approaches promise to be able to identify autism from voice only. These models underestimate how diverse the contexts in which we speak are, how diverse the languages used are and how diverse autistic voices are. Machine learning approaches need to be more careful in defining their limits and generalizability.

## Introduction

Voice atypicalities are argued to be markers of autism across verbal, pre-verbal and nonverbal individuals, albeit with more studies investigating the first group (McCann & Peppe, 2003; Fusaroli et al., 2017). Several machine learning (ML) studies have accordingly attempted to identify a clear acoustic profile of autism, with promising results for supporting assessment processes (Hegde et al., 2019; Mohanta et al., 2020; Verde et al., 2018). However, any potential clinical application requires a careful assessment of how well the ML algorithms generalize not only to new participants from the same study sample, but also to new contexts, subpopulations and languages (Chekroud, 2018; Fusaroli et al., 2017; Low et al., 2020; Rocca & Yarkoni, 2020; Vandenbroucke et al., 2007; Yarkoni, 2020). The assessment of generalizability of ML models of vocal markers of autism is still largely missing, and is the focus of this study.

Currently, clinical features of autism are predominantly assessed using clinical expertise, along with standardized protocols such as those in Autism Diagnostic Interview-Revised (ADI-R) and Autism Diagnostic Observation Schedule-Generic (ADOS-G), which are commonly used in research practice, see (Lord et al., 2008). The process is time-consuming and resource-intensive, as it relies heavily on high-quality training to ensure the validity of the examiners ‘ ratings (Drimalla et al., 2020). Finding more automatically measurable markers could support, simplify and increase the reliability of the assessment process.

Vocalization patterns are associated with diverse cognitive and motor abilities, as well as emotional states and levels of stress (Talkar et al., 2020; Williamson et al., 2017). Clinical features of autism include repetitive behaviors, socialization and communication atypicalities, cognitive deficits, anxiety and sensory overload (Benson & Fletcher-Watson, 2011; Scheerer et al., 2020; Vargason et al., 2020), which are consequently likely to be reflected in autistic voice patterns (e.g., cognitive deficits resulting in longer or more frequent pauses; or anxiety resulting in increased acoustic jitter). Therefore, atypical vocalization patterns could serve as markers for these characteristics (Vargason et al., 2020; Yankowitz et al., 2019). While it is well accepted that autistic individuals often have atypical voices, e.g. sing-songy or monotone intonation and what has been referred to as “inappropriate” prosody (Baltaxe & Simmons, 1985; McCann & Peppé, 2003; Patel et al., 2019), acoustic investigations of the physical properties of voice underlying such perceptions often present weak or inconsistent findings. A recent meta-analysis identified increased and more variable fundamental frequency, as well as longer and more frequent pauses as common characteristics (Fusaroli et al., 2017). These differences were, however, small and could only partially be replicated on new samples, across biological sexes, and languages (Fusaroli et al., 2018, 2022). Several ML studies have tried complementary approaches to these piecemeal approaches. Because ML approaches can explore large numbers of features at once, they can make use of the many acoustic features made available by modern speech processing techniques that would be too numerous to deal with in traditional statistical frameworks. Further, recent developments in neural networks allow the identification of the most relevant features to the task at hand from the raw speech signal (Badshah et al., 2017; Schneider et al., 2019; Sechidis et al., 2021). Capitalizing on this, ML models have been shown to be able to accurately differentiate previously unheard voice samples from new autistic and neurotypical (NT) individuals relying on vocal characteristics alone, with correct classification ranging from 57 % to 96 % of the samples (Fusaroli et al., 2017).

However, ML algorithms are so apt at finding patterns that they often overfit to the data and learn distinctive patterns that are not present in datasets from other studies (Chekroud, 2018; Kuhn & Johnson, 2013). For instance, one study (Bone et al., 2013) showed that algorithms with high performance in identifying autistic children in *The Interspeech 2013 Autism SubChallenge* dataset were relying on the different background noise and reverberation present in those specific recordings, more than on the actual voices. In other words, were the recording contexts to be switched between autistic and neurotypical children, or new recording contexts being used (e.g., the test being performed in a different school), the algorithms would likely perform at chance level. This example illustrates the importance of ensuring the generalizability of machine learning models: the ability of the algorithms to perform accurately across different datasets. Generalizability has an impact on the inferences one can draw from a study. For instance, it can indicate whether the patterns found are related to an underlying biological atypicality in autism (such as differences in motor control of the vocal cords) across all contexts and languages, or whether they are more specific to e.g., situations of high social pressure, such as a conversation with a stranger. Generalizability has also an obvious impact on clinical applications, which require a clear delineation of how well algorithms might generalize to different recording contexts, subpopulations, and tasks before one can even consider real-world use. One should not only know whether an algorithm can be used as-is in a new context or with a new microphone, but also be aware of biases in assessments, e.g., of specific socio-demographic groups, which might have strong ethical and practical implications (Rocca & Yarkoni, 2021; Varoquaux & Cheplygina, 2021).

Little work has been done to examine the generalizability of the performance of ML models on vocal markers of neuropsychiatric conditions, including autism (Fusaroli et al., 2017; Low et al., 2020; Parola, Simonsen, Bliksted, Zhou, et al., 2020). Algorithms might provide a reliable performance (i.e., sensitivity and specificity in classifying data) only if the data keep certain properties constant: e.g., recording with the same type of microphone, minimizing movement in the speaker relative to the microphone, using a certain task for elicitation, or only assessing native speakers of a given language and possibly sociolect. These limitations by no means spell the end for ML as a diagnostic tool, but it is important to acknowledge such potential model limitations to avoid inappropriate interpretation of the results, or overselling of practical applications.

This study specifically aimed to investigate the generalizability of ML models of vocal markers of autism. We searched the literature to identify a promising ML model and replicated its training - within a highly conservative pipeline - on a 3-study, cross-linguistic dataset. This allowed us to investigate the following three questions:

- Q1: How well do models generalize to different participants from the same study, performing the same task and speaking the same language?
- Q2: How well do models generalize to the same participants performing a different task (describing videos vs. repeating a story) in the same language?
- Q3: How well do models generalize to participants from a different study, performing the same kind of task (repeating a story) but speaking a different native language?

## Methods

### Model identification

To identify promising algorithms, we updated the literature search for ML studies of vocal markers of autism reported in Fusaroli et al (2017), identifying 23 studies (for a more elaborate description of the procedure and a table summarizing the studies, see Appendix A1, Table S1). We then selected one of the several algorithms that could be applied on relatively small samples with standard computational setups (e.g., trained Support Vector Machine (SVM) or random forest algorithms as opposed to deep neural networks), and transparently reported the methodological choices. We focused on relatively simple models, as they tend to be less prone to overfitting, therefore, if we were to find generalizability concerns (i.e., inability to accurately identify autistic participants in new samples), such concerns would be even more relevant for models like neural networks, which are more complex and more likely to overfit. For our methodological model we were mainly inspired by Shahin et al. (2019), a study based on SVM and rigorous cross-validation (CV), reaching high performance in predicting autism from voice. The authors reported an accuracy of 0.88, that is, 88% of the samples were correctly classified, and an F1 score of 0.90. F1 scores are used to calculate performance when e.g., the groups to be identified have a different number of cases, and is defined as the harmonic mean of precision (proportion of true positive examples among the examples that the model classified as positive) and recall (proportion of correctly identified positive examples over total positive examples):

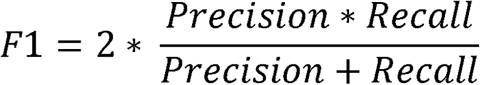

SVM is often used for binary classification problems, and has been shown to perform better than e.g. neural networks and Naive Bayes when there is a limited amount of training data and a high number of features (Kirk, 2017), which is the case in the current study.

Note that different setups relying on more advanced speech recognition technologies (e.g., Hidden Markov models, or recurrent neural networks) applied to large datasets (i.e., thousands of hours of recordings) could be more promising in achieving high levels of performance. However, the amount of data required is unrealistic at this stage, and these tools are currently not commonly employed for the identification of vocal markers of autism.

### Pipeline

This study sets the methodological choices from Shahin et al. (2019) in a highly conservative and fully reported pipeline, which relies on cross-validated training procedures and held-out testing sets, that is, it ensures the model is trained (fitted) on one subset of the data (the training dataset), while its performance is assessed on a different subset of the data (the held-out dataset). Figure 1 provides an overview of the pipeline, which is discussed in detail below.

**Figure 1.**
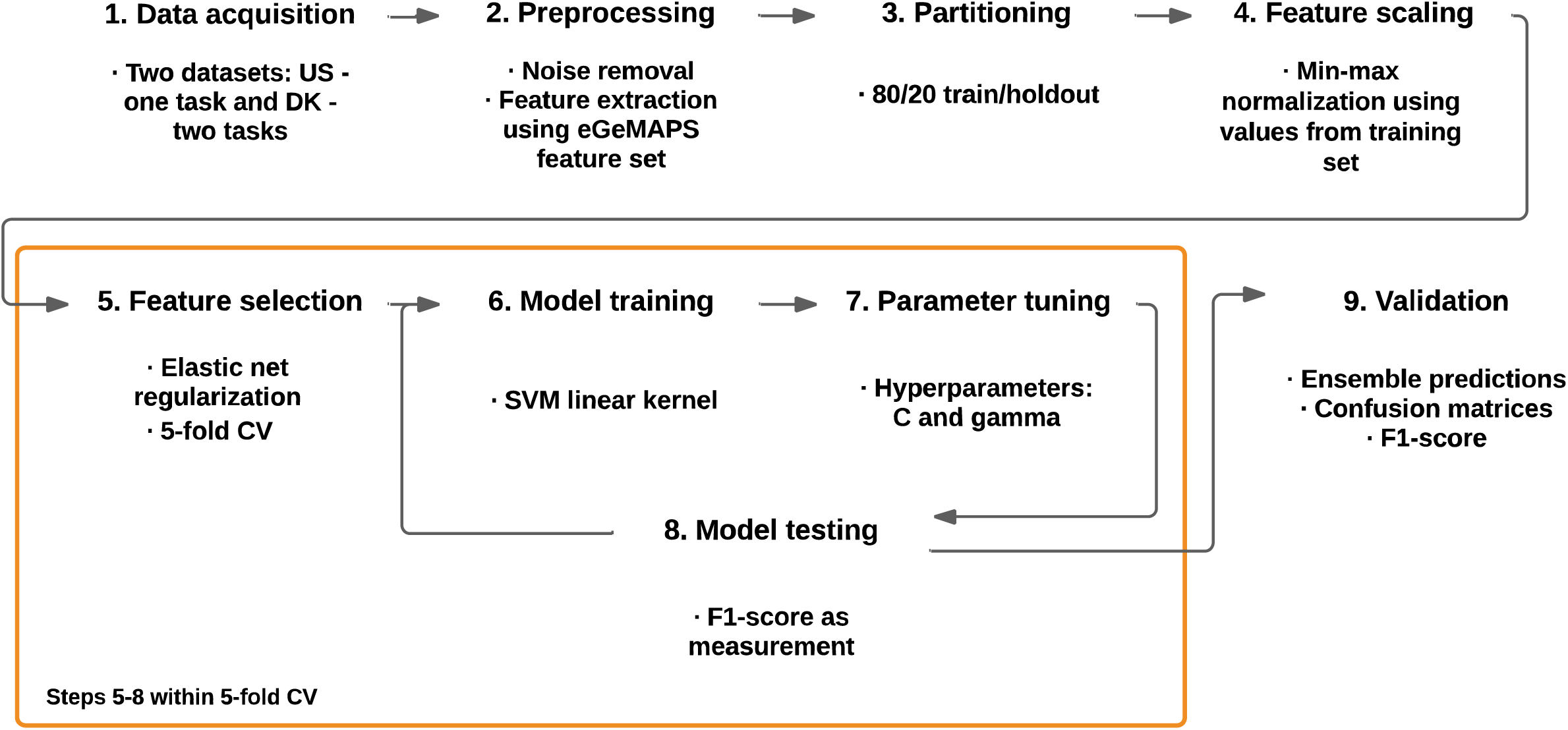
Overview of the machine learning pipeline. Purple refers to the general steps whereas green refers to specifics. “US” indicates the US English study, “DK” the two Danish studies. eGeMAPS indicates the Extended Geneva Minimalistic Acoustic Parameter Set (Eyben et al 2016). Train and holdout indicate respectively the portions of the dataset on which the model is fitted (trained) and tested, respectively. Elastic net indicates a regularization procedure used to reduce the number of features included in the model. SVM RBF indicates the model fitted: Support Vector Machine with a Radial Basis Function. C and gamma hyper-parameters indicate how much error is tolerated on the margin and how curved the margin is expected to be, respectively.

### Data sources

The dataset used in this study consists of voice recordings collected in previous studies for other purposes and their content has been analyzed in other published research (Cantio et al., 2016; Fusaroli et al., 2022; Grossman et al., 2013). For demographic, clinical and cognitive information about the participants, see Table 1^1^.

**Table 1.**
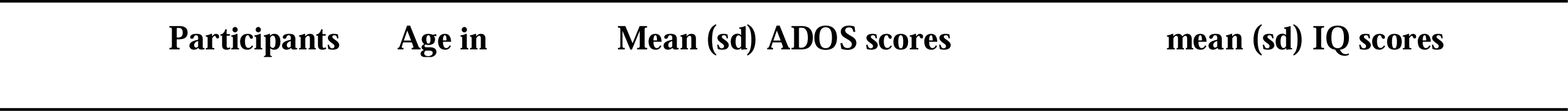
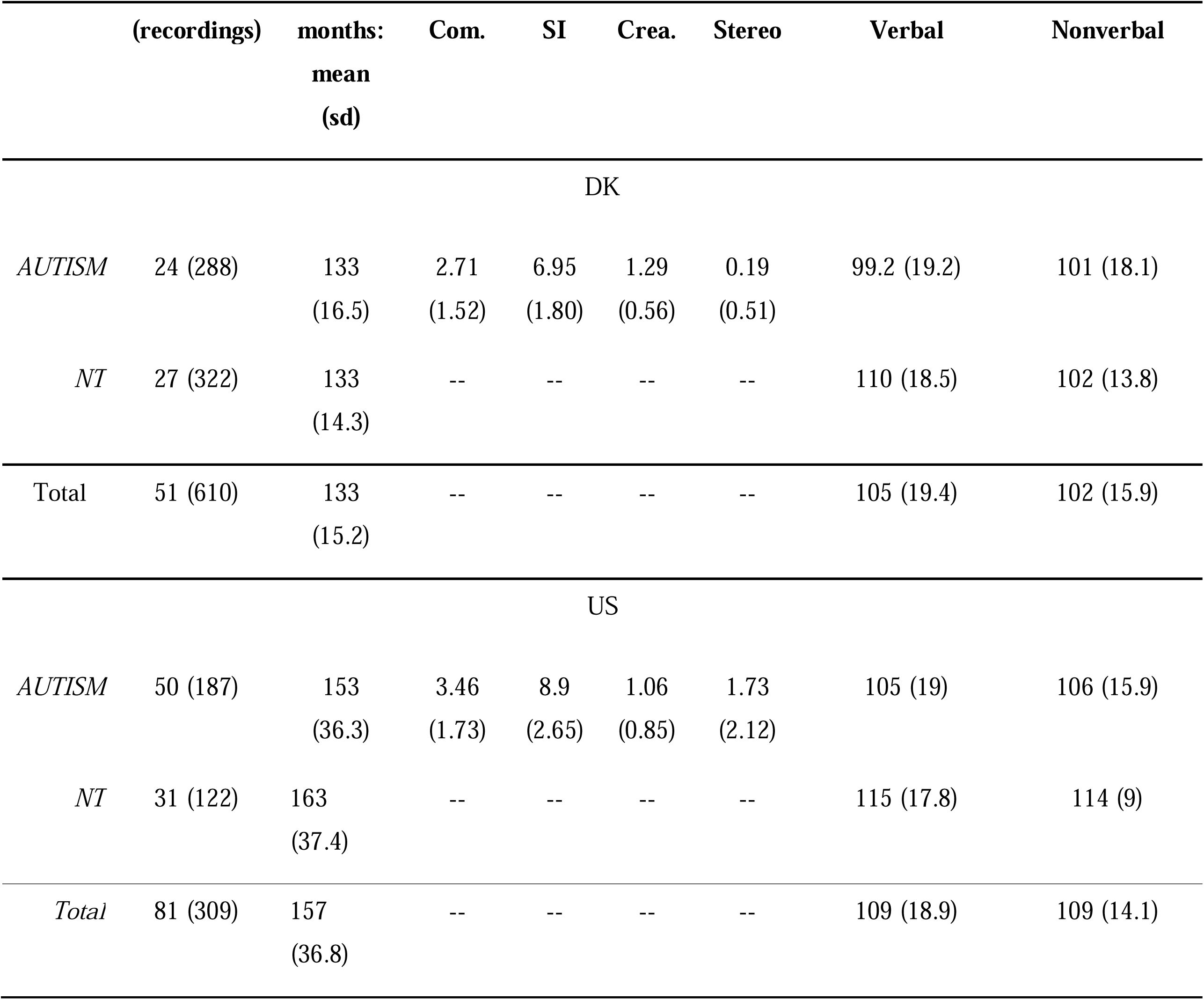
Demographic and clinical features of the participants. DK stands for Danish samples, while US for US English samples. ADOS subscores are abbreviated: “Com.” stands for Communication, “SI” for reciprocal Social Interaction, “Crea.” for Creativity and “Stereo” for Stereotyped Repetitive Behaviors. IQ stands for intelligence quotient.

|  | (recordings) | months:<br>mean<br>(sd) | Com. | SI | Crea. | Stereo | Verbal | Nonverbal |
| --- | --- | --- | --- | --- | --- | --- | --- | --- |
| DK |  |  |  |  |  |  |  |  |
| *AUTISM* | 24 (288) | 133<br>(16.5) | 2.71<br>(1.52) | 6.95<br>(1.80) | 1.29<br>(0.56) | 0.19<br>(0.51) | 99.2 (19.2) | 101 (18.1) |
| *NT* | 27 (322) | 133<br>(14.3) | -- | -- | -- | -- | 110 (18.5) | 102 (13.8) |
| Total | 51 (610) | 133<br>(15.2) | -- | -- | -- | -- | 105 (19.4) | 102 (15.9) |
| US |  |  |  |  |  |  |  |  |
| *AUTISM* | 50 (187) | 153<br>(36.3) | 3.46<br>(1.73) | 8.9<br>(2.65) | 1.06<br>(0.85) | 1.73<br>(2.12) | 105 (19) | 106 (15.9) |
| *NT* | 31 (122) | 163<br>(37.4) | -- | -- | -- | -- | 115 (17.8) | 114 (9) |
| Total | 81 (309) | 157<br>(36.8) | -- | -- | -- | -- | 109 (18.9) | 109 (14.1) |

We collected two Danish and US English datasets involving 74 autistic participants and 58 neurotypical (NT) participants, all with verbal and non-verbal cognitive function within the typical range. Initial data collection was approved by the relevant local IRB ‘s (at Emerson College for the US data, at Region Syd for the Danish data), and the current analysis was deemed exempt from the need of ethical approval by the ethical committee at Region Midt. Each participant recorded several audios, for a total of 919 unique recordings approximately circa 5 hours and 45 minutes in total. The Danish dataset included 24 autistic participants (288 recordings, 88 minutes) and 27 NT participants (322 recordings, 107 minutes), retelling stories (Memory for stories, Reynolds & Voress, 2007) and freely describing short videos (Abell et al., 2000). The US English dataset included 50 autistic (187 recordings, 149 minutes) and 31 NT (122 recordings, 56 minutes) participants, retelling stories (Grossman et al., 2013). The recordings had been collected and analyzed for different purposes in previous studies (Cantio et al., 2016; Fusaroli et al., 2022; Grossman et al., 2013).

Note that the two samples are only roughly matched. While cognitive function and clinical features as measured by ADOS are largely overlapping, US participants are a bit older and varied in age than Danish ones. Further, the language spoken, while always a Germanic one, is obviously different. In particular, Danish is often characterized as an atypical language, with strong reduction in consonant pronunciation (Trecca et al., 2021), although to our knowledge no systematic comparison with US English has been performed. These differences are not a conceptual problem for the present analyses: it is crucial to assess whether so-called vocal markers of autism can generalize across corpora with different characteristics and explore how these differences might matter for the generalizability of the findings.

### Preprocessing

The recording equipment and procedure were consistent within language, but not further specified in the original studies. Experimenter voice, background room noise, reverberation, and hum were removed from the audio recordings using iZotope RX 6 Elements (Izotope, 2017). Long-term average spectra for each recording were inspected for possible noise artefacts and further cleaned if any were found (Olsen, 2018). Following Shahin et al. (2019), we extracted the standardized voice feature set eGeMAPS with the open-source software OpenSmile (Eyben, 2015, p. 201; Eyben et al., 2010, 2016). The feature set contains 87 features, which are described in Appendix A2, Table S2. Note that there are concerns about the reliability of acoustic features over time and across noisy environments (see e.g., Stegmann et al., 2020). This is an important concern that should be further investigated. Given that the previous literature on vocal markers of autism acritically relies on acoustic features like those extracted by OpenSmile and that we aim at assessing the generalizability of findings from the previous literature, we chose not to further select features for their reliability. Nevertheless, to mitigate the issue we relied on summary features (median and interquartile range) across each recording and on repeated measures for each participant (2-10 recordings per participant per task). Note that by using summary features of this kind, we are neglecting temporal variation, which are likely to be relevant as markers of autism (Bone et al., 2014; Fusaroli et al., 2015). However, our goal is to assess the generalizability of current ML practices in the field and we leave the identification of better features than those currently used to future work.

All features in the training data were scaled using min-max normalization to achieve a dataset with a common scale without losing information or distorting differences in the range of values (Singh & Singh, 2020). The values used to normalize the training set were saved and used to normalize the test set, thus avoiding potential information leakage between the two sets.

### Model training procedure

To investigate Q1, Q2 and Q3, three models were trained. The first model was trained on the Danish Animated Triangle data, the second on the Danish storytelling data, and the third on the US storytelling data. After training, the models were tested on different test sets. The combinations of training and test sets can be seen in Figure 2.

**Figure 2.**
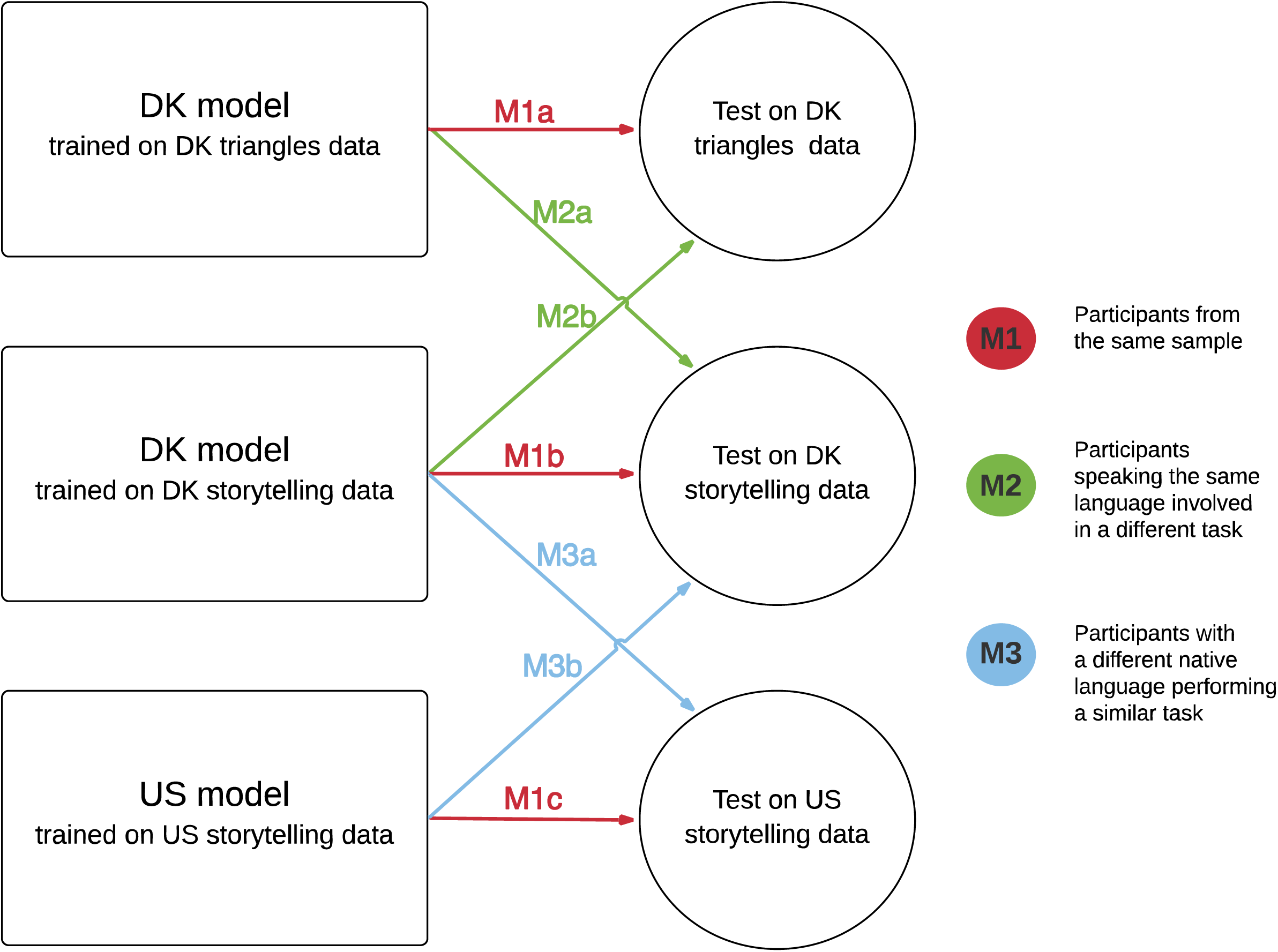
Overview of model training and testing. Each square indicates a 5-model voting ensemble trained on one dataset. Each circle indicates a held-out testing dataset. The arrows show which models are tested on which held-out dataset and their color-codes for the question that is being asked: Red for Q1 (new participants from the same sample, separated in M1a, M1b, and M1c according to the datasets involved), green for Q2 (same participants, different task, separated in M2a, and M2b according to the datasets involved), and blue for Q3 (same task, different language, separated in M3a, and M3b according to the datasets involved). See also Table 2.

Each dataset was partitioned into a training set and a test set with an 80/20 split, stratified by participant ID; all data points for a given participant ended up in the same partition. We also ensured the test set included an equal number of participants from each diagnostic group (total number of voice recordings differed in some cases - see Appendix A5, Table S10) to support more robust out-of-sample performance. This test set was used to answer questions about the generalizability to different participants from the same sample, performing the same task (Q1). Unfortunately, using a test set in this condition limits the amount of datapoints we can calculate performance on, with a lower limit for Q1b of 18 participants and 36 recordings, which is still comparable to or better than the studies reviewed in Fusaroli et al (2017). When testing Q2 and Q3, the test dataset consisted of either the same participants performing a different task (Q2) or participants with a different native language performing the same kind of task (Q3).

To prevent overfitting as much as possible, we relied on a strict five-fold cross-validation of the training process. In other words, the training dataset was split into five folds (roughly equally sized subsets) balanced by participant ID, so that recordings from the same participant only appeared within a single fold, and the testing fold only contained never-before-seen participants. Each of these five folds was then used as a validation set for a training set composed of the other four folds (see figure 1). Within each of these five CV training sets, we performed feature selection, and tuned y (gamma, or the curvature of the margin between classes) - and C (or the error tolerated at the margin) – hyper-parameters using a grid search method (see Appendix A3 for details). SVM classifiers were implemented with radial basis function (RBF) kernels (Pedregosa et al., 2011; Van Rossum & Drake, 2009). Due to the cross-validation process, this yielded five trained SVM models for each of the three datasets used.

### Majority voting

The 5 SVM models were used to assess the test sets, and their predictions were combined into a single voting ensemble (Brownlee, 2020; Hansen et al., 2021; Sechidis et al., 2021). Each model made a prediction for each voice recording in the test set, and the ensemble model gave a final predicted class based on the majority of these model votes. Note that other systems beyond majority rules have been developed, e.g., Mixture of Experts with weights based on similarity between test and training data (Hansen et al., 2021; Sechidis et al., 2021). Combining or utilizing multiple models within a single model – such as an ensemble model – benefits performance and generalizability, since no two models are likely to overfit in the same way and different models can compensate for each other ‘s biases (Buracas & Albright, 1993; Hong & Page, 2004; Tang et al., 2005).

### Impact of amount of data available

Given previously expressed concerns about the limited sample sizes available in studies of vocal markers of autism, we also exploratorily test whether systematically manipulating the amount of training data available affects the algorithms ‘ test performance. We repeated our analyses varying the numbers of folds employed in the cross-validation: 2, 3, 4, and 5. Using a 2-fold cross-validation, the algorithm has access to about half the participants in the dataset as training data. Using a 3-fold cross-validation, to two thirds of the participants; with 4-fold cross validation to three fourth and with 5-fold to four fifths. If data availability - within the limits of our datasets - is a key factor in the generalizability of the patterns found we expect test performance to increase with the number of folds.

### Software implementation and open science

All steps in the analysis – except for the de-noising – are implemented using open software. The feature extraction software toolkit openSMILE 3.0 was used to extract the feature sets GeMAPS, eGeMAPS and ComParE from the speech signals (Eyben et al., 2010). Data cleaning, including removal of outliers and normalization, and feature reduction using ElasticNet was implemented using R (RStudio Team, 2020). Model training for all kernels and SVMs as well as validation and testing was carried out using the Scikit-learn module in Python. The source code for the analysis can be found at: https://osf.io/9mtpk/?view_only=fa4497a6478b48118e6d15b31cc07567

## Results

A crude overview of the performance of the seven models (three testing on the same task in the same language, two testing on a different task in the same language, two testing on the same task in a different language) is given in Table 2. We include plots of precision (proportion of true positive examples among the examples that the model classified as positive) and recall (proportion of correctly identified positive examples over total positive examples) (Figure 3) on test sets to enable a more precise error analysis of the results. Model performance is evaluated using F1-score, the harmonic mean of precision and recall. Confusion matrices displaying the raw performance can be seen in Appendix A4, Tables S3-S9. Receiver Operating Curve plots can be seen in Appendix A4, Figure S1. Data availability – within the limits of our datasets - only minimally impacts generalizability of the patterns. As presented in Figure 4, while 2-fold cross validation seems to lead to worse results, there is no reliable increase in performance between 3- and 5-fold cross-validation.

**Table 2.** Overview of model performance on different test sets, F1-scores. The colors code the question tackled by the specific test set: M1 (red) answers question 1; M2 (green) answers question 2; M3 (blue) answers question 3.

| Tested on →<br>Trained on<br>↓ | DK triangles data | DK storytelling data | US storytelling data |
| --- | --- | --- | --- |
| DK triangles data | **M1a** - Q1<br>0.59 | **M2a** - Q2<br>0.60 | -- |
| DK storytelling data | **M2b** - Q2<br>0.66 | **M1b** - Q1<br>0.89 | **M3a** - Q3<br>0.45 |
| US storytelling data | -- | **M3b** - Q3<br>0.40 | **M1c** - Q1<br>0.64 |

**Figure 3:**
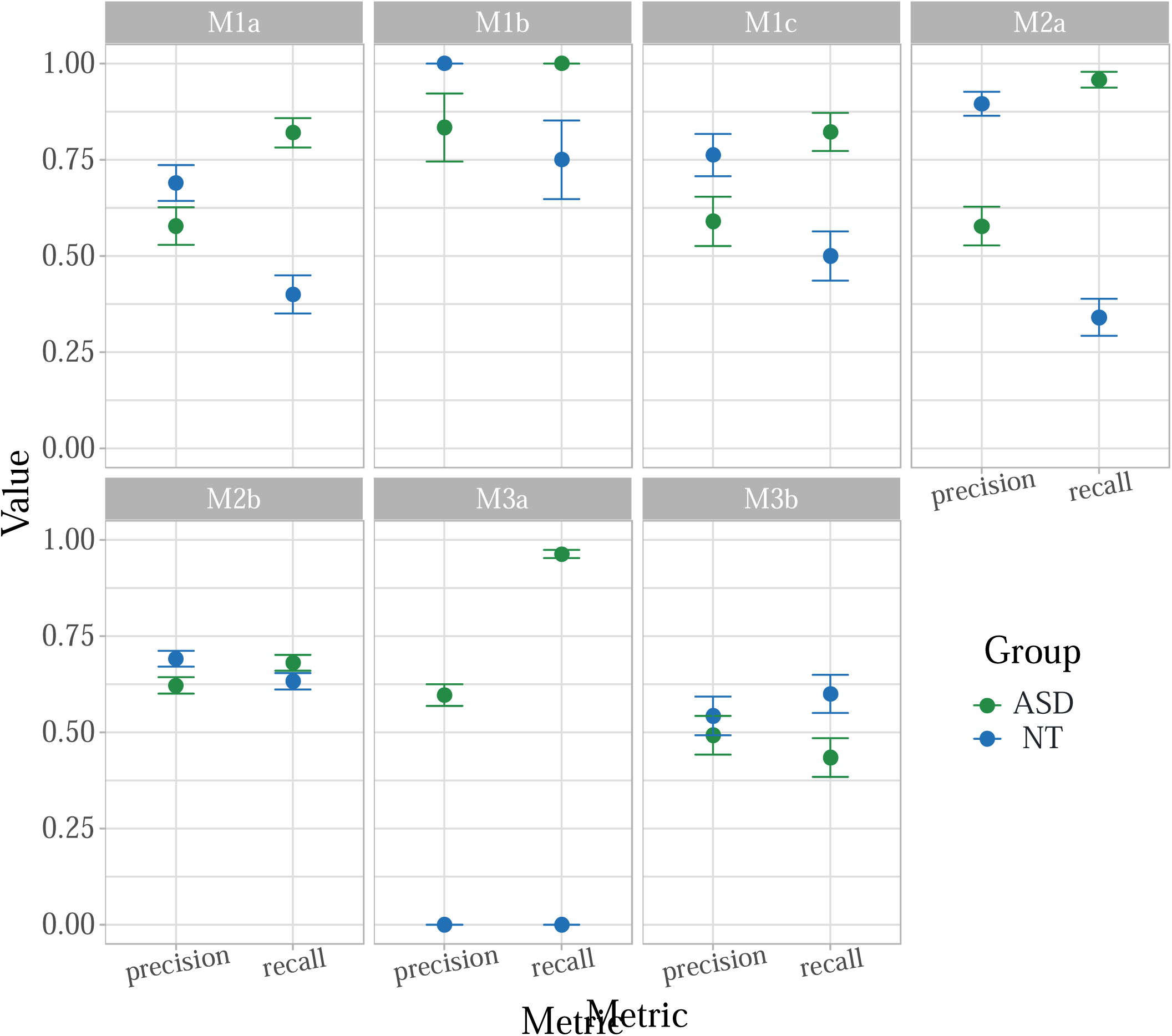
Precision (fraction of true positive examples among the examples that the model classified as positive) and recall (fraction of correctly identified positive examples over the total of actual positive examples) for autistic (ASD) and neurotypical (NT) participants, respectively, for all models. Compare with table 2 (overview of models). To interpret the panels, consider e.g., panel M1a assessing the performance of the algorithm trained on Danish triangle data and evaluated on the test set from the same task and language. Precision (on the left) is higher for neurotypical participants (if the model says “neurotypical”, it is more accurate than if it says “autistic”), while recall is higher for the autistic group (autistic participants are more likely to be correctly identified, than neurotypical ones).

**Figure 4.**
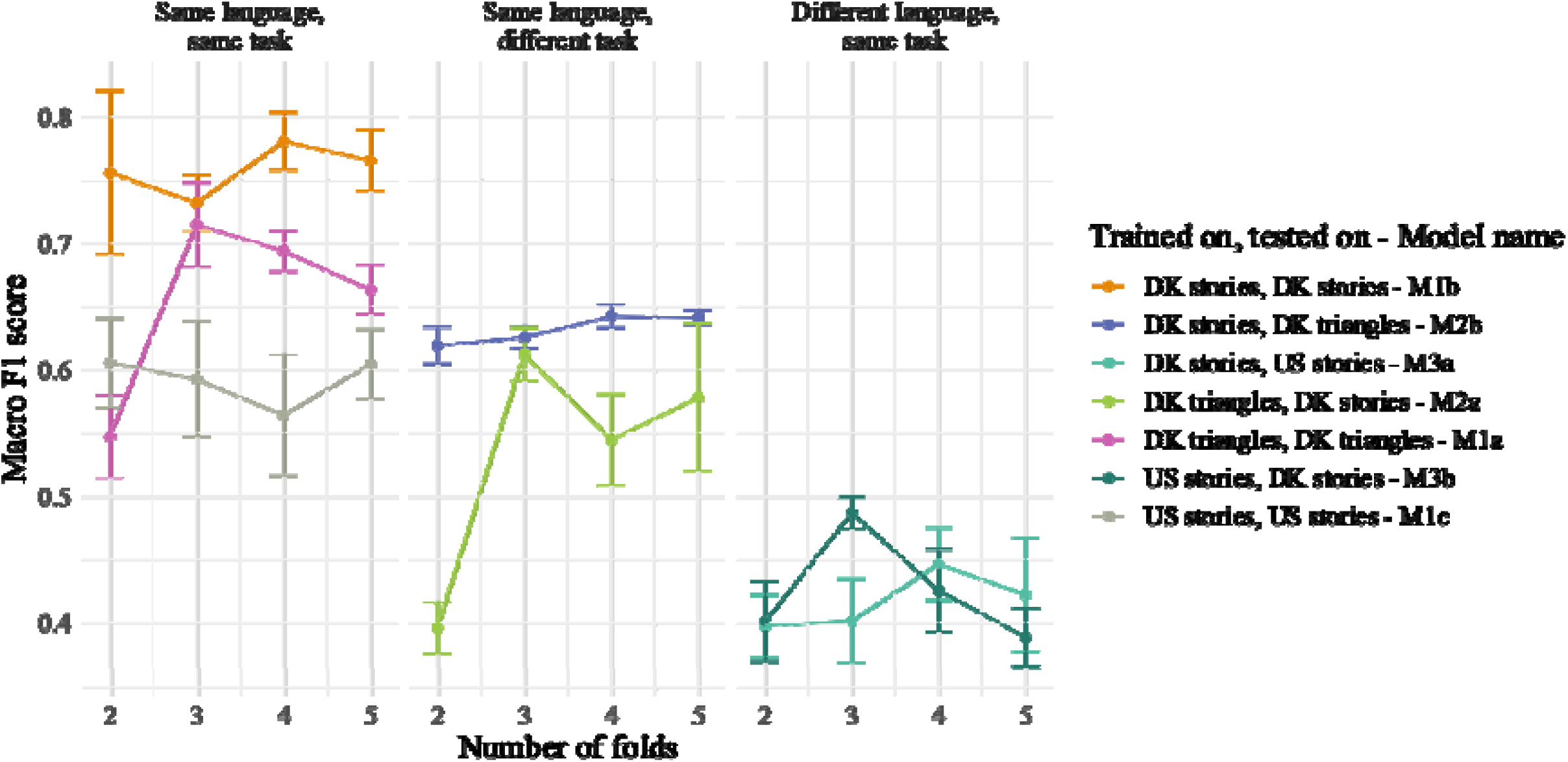
Test performance as a function of data availability. We repeated all analyses using 2, 3, 4, or 5 folds in the cross-validation process, thus systematically manipulating the amount of training data available to the classifiers (fewer folds imply less training data) and assessed their test performance. Error bars indicate standard errors across 5 runs of the algorithms.

## Discussion

In this work we systematically assessed the generalizability of ML results on vocal markers of autism. We relied on cross-linguistic data to assess whether the results would generalize: (Q1) to different participants from the same study, performing the same task; (Q2) to a different task performed by the same participants; and (Q3) to participants from a different study, with a different native language performing the same type of task. Below, we briefly discuss our specific findings in relation to these three questions, and the specific features of our dataset. We then advance more general recommendations for how future ML studies could take generalizability concerns more seriously.

When assessing Q1, we found that all models, independently of the dataset they were trained on, were above chance at identifying autistic participants amongst new participants from the same study, with performance within the range of previously published articles: accuracies between 62% and 87% (Fusaroli et al., 2017). However, the very same training procedure yielded highly variable results on the three datasets (F1-scores between 0.59 and 0.89), with the lower scores being of dubious usefulness in clinical contexts. Previous studies suggest that the different demands of the diverse tasks used to collect speech recordings impact how clearly vocal atypicalities are expressed by participants (Fusaroli et al., 2017; Parola, Simonsen, Bliksted, & Fusaroli, 2020). In particular, more cognitively and socially demanding tasks tend to show bigger vocal differences between neurodiverse and neurotypical populations. However, we doubt that this is the explanation for our findings, since we do not see any reason to suggest that retelling stories in Danish is inherently more challenging than describing previously watched videos in Danish, or retelling stories in US English. Additionally, exploratory analyses on the very few female participants suggested little ability to generalize to new participants of another biological sex. These findings are worrying, since they indicate that even re-training ML models on one ‘s own dataset is not guaranteed to yield the same performance as published findings. One potential explanation for this is the relatively small size and homogeneity of the datasets involved both in previous studies and in the current investigation. While manipulating sample sizes within the limits of our datasets did not yield reliable changes in performance, employing a much larger sample size with increased attention to represent the heterogeneity of the autistic population might yield more stable results.

When assessing Q2, we found again that the models were able to assess the same participants performing a new task with an accuracy above chance, but generally lower than on the original task (F1 scores of 0.60 and 0.66) and of dubious usefulness for any practical concern. The participants are the same, the tasks are relatively similar, and the conditions (and date) of recording are exactly the same, so we speculate that some overfitting to the specific task is at stake.

When assessing Q3, we found that the models were not able to assess other languages with the same type of task, with performance dropping below chance (F1 scores of 0.4 and 0.45). Exploration of the performance (e.g., Table S8) indicates that in some cases all autistic participants are misclassified, and therefore showing that the algorithm drastically fails at the task. An exploratory analysis focusing only on age matched individuals across the two languages yielded qualitatively identical results (lack of any generalization of findings). Relatedly, one previous study relied on a cross-linguistic corpus (English, Hebrew, Swedish and French, with no reported background noise removal procedure) reported inconsistent results and low performance as to whether new participants speaking a different language could be identified above chance (Schmitt et al., 2016). Transferring a ML model of autistic voice across languages is thus non-trivial, and we should not expect findings to generalize without re-training the model.

These findings raise strong concerns as to the generalizability of ML models of voice in autism (for similar worries in the context of other neuropsychiatric conditions, see Arora et al., 2019; Berisha et al., 2021; Stegmann et al., 2020). First, performance of the same model trained on relatively similar datasets changed quite drastically, indicating that evaluations of ML models on one dataset might not be representative of the results that could be achieved on a different dataset. Second, generalizability across relatively similar tasks (description of a narrative video vs. retelling of a story) is low and of dubious usefulness, even when keeping participants and native language constant. Third, even when testing phylogenetically close languages (US English and Danish being both Germanic languages), the acoustic patterns found by our models could not generalize. One could speculate that testing on a Romance language (e.g., Italian), or a non-Indo-European language (e.g., Mandarin Chinese) would compromise generalizability even further.

In summary, while models can be trained on novel datasets to reach roughly comparable performance to original reports, there is considerable variance, and these levels of performance do not generalize beyond the specific task and language. Since these issues emerge even with relatively simple SVM classifiers, with a limited amount of hyperparameters to set, we expect them to be even more prominent in more complex model architectures such as convolutional neural networks or other deep learning methods, as these models are known to be more prone to overfitting (Kuhn & Johnson, 2019). These issues are further exacerbated by several factors: the study of autistic voice is generally constrained by limited data availability (Fusaroli et al., 2017) and it is focused on clear-cut autism vs neurotypical comparison groups, instead of more realistic situations including comorbidities and non-autism-related neurodevelopmental conditions in both groups. We therefore argue that it is crucial to explicitly tackle the problem of generalizability in order to advance the field.

Our study relied on samples in which all participants were children, exclusively boys, and had normal range verbal and cognitive functions. One could question the relevance of an autism assessment for children as old as those in our age range. The goal of this study was a more general assessment of the current ML attempts at identifying behavioral – and vocal in particular - markers of autism, which means the specific age range is not in focus and we expect results – how well models generalize – to extend to studies of other age groups. Further, the identification of vocal markers of autism could lead to the support of the assessment of clinical, cognitive and social features as they develop, which is an important issue in the current age range. Being limited to boys, the study relies on a more homogeneous set of acoustic features, which might have increased performance. Given the failure to generalize the findings, we do not think this an issue for our conclusions, but we do advocate for more work to be done beyond male voices in autism. Finally, the relatively high range of verbal and cognitive abilities represented in the samples also questions how representative our findings are of the full autistic spectrum. While this is certainly a limitation and more work should be dedicated to non- or minimally verbal vocal production, it is also a strength. Autistic and neurotypical participants both produce verbal content, which can be more evenly compared in terms of its acoustic properties, then in presence of no production or non-verbal vocalizations. More research will be needed to extend this line of research to preverbal and non-verbal individuals.

Moving beyond the specific case of our own datasets and ML procedures, we want to make a more general perspective for taking generalizability concerns more seriously in ML research, and provide the following three key guidelines that we elaborate on below:

1. Explicitly discuss generalizability concerns (as related to the question at hand)
2. Explicitly assess relevant generalizability of performance
3. Consider the use of multi-datasets ML techniques

### Explicitly consider generalizability concerns as related to the question at hand

In a recent review, Low et al. (2020) emphasize that many studies of voice in psychiatric conditions do not measure out-of-sample performance, e.g., via cross-validation or testing on hold-out subsets of the datasets. Our findings indicate that even when correctly applied these forms of out-of-sample validation of ML models might not provide reliable measures of generalizable performance, depending on the research goals and application needs. For instance, a study trying to identify whether there is an acoustic profile of autism in general should explicitly discuss that this means the findings should generalize across biological sex, language, socio-demographic and ethnic groups, but also that it does not need to generalize across recording settings, given the research question at stake. On the contrary, a study trying to provide tools for the assessment of autism via voice markers across a variety of clinics and contexts should also consider varying recording setups (e.g., microphones, background noise, reverberation differences), but might have lower cross-linguistic generalizability needs (depending on the context of use). Needless to say, requirements will often be domain-specific, but explicitly stating them is a prerequisite for proper usage and cumulative knowledge building within the field, and there has been at least one attempt at establishing such a practice for more traditional statistical approaches (Simons et al., 2017).

### Explicitly assess relevant generalizability of performance

Additionally, we argue that generalizability of the algorithm must not only be discussed, but also thoroughly tested and documented. An explicit assessment of generalizability eases the understanding of a model ‘s capabilities as well as its limitations within the intended use. A first step could be an error analysis, as assessing which errors are made by the model might highlight specific biases, e.g. poor ability to identify autistic women, or participants from specific social and ethnic groups (Achenie et al., 2019). However, that is not enough. As we have shown, relying on data from one study and within-study out-of-sample validation is not a reliable way of ensuring generalizable findings, or even documenting limits of the algorithms and data. This, of course, requires the construction or availability of multiple datasets. The increase in collaboration across labs, construction of consortia such as EU-AIMS, ManyBabies, and Psychological Science Accelerator and the increased spread of responsible open science practices might help with this (Bergmann et al., 2016; Murphy & Spooren, 2012).

### Consider the use of multi-dataset ML techniques

As data from multiple studies become available, new techniques become possible, moving from single study training to multi-study approaches, where models are trained on data collected under diverse settings. A basic approach is simply to train the models on multiple datasets, to better account for the heterogeneity between individuals, contexts, tasks and languages. However, more nuanced approaches may provide more modular and flexible ways of taking advantage of multiple datasets. One such approach is the use of mixture-of-experts (MoE) models. In a MoE-model, separate models are trained, each on a different dataset. When evaluating test cases, each model provides both a prediction, and an evaluation of how similar the single case is to the training set (similarity score). The predictions of each expert are then combined into a final prediction, i.e. by weighting the predictions of the experts proportionally to their similarity score (Sechidis et al., 2021). Thus, if one seeks to construct a model that generalizes to three languages, one could train a model on each language separately and combine their predictions in a MoE-model. If a new language were to become available, a new model could be trained only on the new data and added to the mixture. This approach has already shown promising results in related fields, e.g. in predicting the emotional content of Parkinson ‘s and depressive speech in languages on which the models had not been trained (Hansen et al., 2021; Sechidis et al., 2021). Analogously, modular approaches have been developed for language models, including novel techniques to mix, re-weight, add or entirely remove “experts” according to the task at hand (Gururangan et al., 2021). These models can indeed generalize and adapt to new domains and achieve high performance scores when taking advantage of domain-specific expertise. Another promising line of research is transfer learning, where complex models can be trained on large and more easily accessible datasets of non-clinical speech, and then only re-trained on the smaller clinical datasets; or across clinical datasets (Vásquez-Correa et al., 2019). Developing such techniques within the field of voice-based classification of autism is a highly promising venue to foster higher generalizability.

Finally, we emphasize the importance of sharing openly and fully the modeling process - e.g., strategies to choose hyperparameters, and actual parameter values - to better enable collective investigation of generalizability issues. Accordingly, our choices are fully described in the manuscript and appendices and the code is available on the OSF repository connected to this paper (https://osf.io/9mtpk/?view_only=fa4497a6478b48118e6d15b31cc07567). More research is needed to assess the generalizability of our findings to new data, e.g., including different age ranges, languages, linguistic and cognitive function, co-morbidities and other neurodevelopmental conditions. By sharing our data and code, we hope to make this easier.

### Limitations of current approaches

Our first two recommendations are meant to make issues of generalizability and their conceptual implications more transparent. In this study, we show how implementing them highlights the lack of generalizability of current ML approaches to vocal markers of autism. However, these approaches are likely limited and should be improved to increase the findings ‘ generalizability. In particular, relying on the history of speech recognition techniques and other machine learning applications, we can identify three key limitations in the machine learning approach we assessed: samples, features and algorithms. Our corpora consist of less than 6 hours of recordings from 132 individuals. Albeit equivalent to or larger than most corpora currently used to identify vocal markers of autism, it does not match up to the > 10.000 hours corpora used to train speech recognition algorithms. Vocal production is highly variable, within recording, across repeated recordings and between individuals; a variability that we can just imagine being even larger in autism, due to the heterogeneity encompassed by the condition. Attempting to capture generalizable patterns in small samples is probably of limited usefulness. In this sense, our third recommendation might represent a first step forward by catalyzing the use of multiple datasets capturing more heterogeneity in vocal production. Nevertheless, specific attention to large samples, multiple languages, repeated measures from multiple individuals, and effort in representing a broader portion of the condition is needed. Given the difficulties in collecting such data, large-scale community efforts, shared open data where possible, and cumulative approaches are needed.

A second important limitation is the features extracted. Summary features neglecting the temporal dynamics of vocal productions are likely to neglect relevant aspects of autistic voices. While hand-engineering of temporal acoustic features is certainly an option (Fusaroli et al., 2015), recent developments in time-sensitive algorithms – such as recurrent neural networks dealing with the sequence of acoustic values in time - might also be employed. More radically, in the presence of larger datasets, deep learning algorithms have also been successfully used to automatically infer relevant features from the data. Such algorithms, however, have their own downsides (e.g., a very strong tendency to overfit to the data, or use features that are irrelevant to the question at stake, such as the structure of background noise). Therefore, more work is needed to assess their generalizability in assessing vocal markers of autism.

## Conclusion

This work investigated the generalizability of ML approaches to autistic voices. While promising algorithms could be retrained on new datasets with performance above chance, variability in performance across algorithms trained on different studies was substantial. Further, the models did not generalize well to different but similar tasks and not at all to new languages. We argue that greater emphasis must be placed on the generalizability of machine learning models of autistic voices. We recommend to 1) explicitly discuss generalizability concerns (as related to the question at hand), 2) explicitly assess relevant generalizability of performance and 3) consider the use of multi-study ML techniques.

## Acknowledgments

We are very grateful to the participants in our studies and to the researchers and clinical practitioners who supported the collection of the data we analyzed in this project. We wish to thank our funding sources: the Interacting Minds Centre seed funding “Clinical Voices” and “Clinical voices in the wild”; and the Danish Independent Research Council collective project “The Puzzle of Danish”. Lasse Hansen, Alberto Parola and Kostas Sechidis provided invaluable feedback.

## Footnotes

1 Note that female participants’ data in our original datasets were very sparse: e.g., the US dataset contained only 5 autistic girls (3.8%). Since exploratory analyses showed little ability of the models to generalize to female voices, but female participants were too few and unbalanced across datasets to be able to draw any conclusion, we excluded them from the current study, and therefore the table. For a more detailed analysis of differences in vocal markers of autism depending on biological sex see Fusaroli et al. (2021).

