## Supplementary Materials for "Vocal markers of autism: assessing the generalizability of machine learning models"

### Appendix A1 – Previous Machine Learning Studies

The studies chosen for this work are based on a literature search in Google Scholar following the review by Fusaroli et al. (2017). The search included studies published between 2015 and 2020 in order to only capture studies published later than the review. As this study is concerned with machine learning approaches to voice and autism, the search term "machine learning" was added to the search terms from Fusaroli et al. (2017). As the data in our study was obtained from the age group 8 - 19 years, studies which included the terms “infancy" and “prelinguistic" were excluded. The search terms used were:

(prosody OR intonation OR inflection OR intensity OR pitch OR fundamental OR frequency OR speech OR rate OR voice OR quality OR acoustic) AND autis* AND "machine learning" -infan* -prelingui*.

Along with the machine learning studies identified by Fusaroli et al. (2017), this search yielded 23 relevant studies in total (see Table S1) of which one was found to be particularly relevant due to 1) methodological transparency and 2) theoretical motivation and background. Accordingly, the study by Shahin et al. (2019) was chosen as subject for replication in this study. Shahin et al. (2019) trained models to categorize various Speech Sound Disorders (SSD).

Table S1 – Literature building machine learning models of vocal markers of autism – as identified in our literature search.

| # | Reference |
| --- | --- |
| 1 | Asgari, M., & Shafran, I. (2018). Improvements to harmonic model for extracting better speech features in clinical applications. Computer Speech & Language, 47, 298-313. |
| 2 | Asgari, M., Bayestehtashk, A., & Shafran, I. (2013, August). Robust and accurate features for detecting and diagnosing autism spectrum disorders. In Interspeech (Vol. 2013, p. 191). NIH Public Access. |
| 3 | Bedoya, S., Katz, N. B., Brian, J., O’Shaughnessy, D., & Falk, T. H. (2020). 3. Acoustic and prosodic analysis of vocalizations of 18-month-old toddlers with autism spectrum disorder. In Acoustic Analysis of Pathologies (pp. 93-126). De Gruyter. |
| 4 | Bone, D., Chaspari, T., Audhkhasi, K., Gibson, J., Tsiartas, A., Van Segbroeck, M., ... & Narayanan, S. S. (2013). Classifying language-related developmental disorders from speech cues: the promise and the potential confounds. In INTERSPEECH (pp. 182-186). |
| 5 | Bone, D., Lee, C. C., Black, M. P., Williams, M. E., Lee, S., Levitt, P., & Narayanan, S. (2014). The psychologist as an interlocutor in autism spectrum disorder assessment: Insights from a study of spontaneous prosody. Journal of Speech, Language, and Hearing Research, 57(4), 1162-1177. |
| 6 | Bonneh, Y. S., Levanon, Y., Dean-Pardo, O., Lossos, L., & Adini, Y. (2011). Abnormal speech spectrum and increased pitch variability in young autistic children. Frontiers in human neuroscience, 4, 237. |
| 7 | Cho, S., Liberman, M., Ryant, N., Cola, M., Schultz, R. T., & Parish-Morris, J. Automatic detection of ASD in children using acoustic and text features from brief natural conversations. |
| 8 | Cummins, N., Baird, A., & Schuller, B. W. (2018). Speech analysis for health: Current state-of-the-art and the increasing impact of deep learning. Methods, 151, 41-54. |
| 9 | Deng, J., Cummins, N., Schmitt, M., Qian, K., Ringeval, F., & Schuller, B. (2017, July). Speech-based diagnosis of autism spectrum condition by generative adversarial network representations. In Proceedings of the 2017 International Conference on Digital Health (pp. 53-57). |
| 10 | Kakihara, Y., Takiguchi, T., Ariki, Y., Nakai, Y., Takada, S., & Kakihara, Y. (2015). Investigation of classification using pitch features for children with autism spectrum disorders and typically developing children. American Journal of Signal Processing, 5(1), 1-5. |
| 11 | Kiss, G., Santen, J. P. V., Prud'Hommeaux, E., & Black, L. M. (2012). Quantitative analysis of pitch in speech of children with neurodevelopmental disorders. In Thirteenth Annual Conference of the International Speech Communication Association. |
| 12 | Kokot, M., Petric, F., Cepanec, M., Miklić, D., Bejić, I., & Kovačić, Z. (2018, June). Classification of child vocal behavior for a robot-assisted autism diagnostic protocol. In 2018 26th Mediterranean conference on control and automation (MED) (pp. 1-6). IEEE. |
| 13 | Kosmicki, J. A., Sochat, V., Duda, M., & Wall, D. P. (2015). Searching for a minimal set of behaviors for autism detection through feature selection-based machine learning. Translational psychiatry, 5(2), e514-e514. |
| 14 | Li, M., Tang, D., Zeng, J., Zhou, T., Zhu, H., Chen, B., & Zou, X. (2019). An automated assessment framework for atypical prosody and stereotyped idiosyncratic phrases related to autism spectrum disorder. Computer Speech & Language, 56, 80-94. |
| 15 | Marchi, E., Schuller, B., Baron-Cohen, S., Golan, O., Bölte, S., Arora, P., & Häb-Umbach, R. (2015). Typicality and emotion in the voice of children with autism spectrum condition: Evidence across three languages. In Sixteenth Annual Conference of the International Speech Communication Association. |
| 16 | Marchi, E., Schuller, B., Baron-Cohen, S., Golan, O., Bölte, S., Arora, P., & Häb-Umbach, R. (2015). Typicality and emotion in the voice of children with autism spectrum condition: Evidence across three languages. In Sixteenth Annual Conference of the International Speech Communication Association. |
| 17 | Memari, N., Abdollahi, S., Khodabakhsh, S., Rezaei, S., & Moghbel, M. (2020, July). Speech Analysis with Deep Learning to Determine Speech Therapy for Learning Difficulties. In International Conference on Intelligent and Fuzzy Systems (pp. 1164-1171). Springer, Cham. |
| 18 | Mohanta, A., Mukherjee, P., & Mirtal, V. K. (2020, February). Acoustic Features Characterization of Autism Speech for Automated Detection and Classification. In 2020 National Conference on Communications (NCC) (pp. 1-6). IEEE. |
| 19 | Nakai, Y., Takiguchi, T., Matsui, G., Yamaoka, N., & Takada, S. (2017). Detecting abnormal word utterances in children with autism spectrum disorders: machine-learning-based voice analysis versus speech therapists. Perceptual and motor skills, 124(5), 961-973. |
| 20 | Pahwa, A., Aggarwal, G., & Sharma, A. (2016, April). A machine learning approach for identification & diagnosing features of Neurodevelopmental disorders using speech and spoken sentences. In 2016 International Conference on Computing, Communication and Automation (ICCCA) (pp. 377-382). IEEE. |
| 21 | Schmitt, M., Marchi, E., Ringeval, F., & Schuller, B. (2016, October). Towards cross-lingual automatic diagnosis of autism spectrum condition in children's voices. In Speech Communication; 12. ITG Symposium (pp. 1-5). VDE. |
| 22 | Shahin, M., Ahmed, B., Smith, D. V., Duenser, A., & Epps, J. (2019, October). Automatic Screening Of Children With Speech Sound Disorders Using Paralinguistic Features. In 2019 IEEE 29th International Workshop on Machine Learning for Signal Processing (MLSP) (pp. 1-5). IEEE. |
| 23 | Wijesinghe, A., Samarasinghe, P., Seneviratne, S., Yogarajah, P., & Pulasinghe, K. (2019, October). Machine Learning Based Automated Speech Dialog Analysis Of Autistic Children. In 2019 11th International Conference on Knowledge and Systems Engineering (KSE) (pp. 1-5). IEEE. |

### Appendix A2 – Acoustic Features

Table S2: Overview of features in eGeMAPS

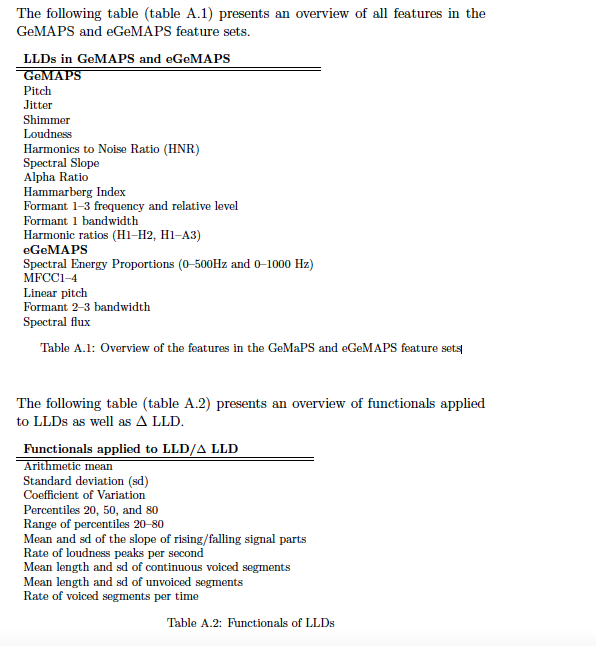

### Appendix A3 – Hyper-parameters

The C- and 𝛾-parameters of the SVM models were chosen using grid search as in Shahin et al. (2019). This means that a grid is constructed for each of the values and the optimal combinations of the two values chosen in a 5-fold CV within the training data. It has been suggested that the distribution of C- and 𝛾-values should not be linear but rather exponential to ensure the same amount of information at each grid point (Pedregosa et al., 2011). Therefore, the grid is constructed from nine C-values with base 0.5 raised to the powers of -1, 0, 1, 2, 3, 4, 5, 6, 7 and eight 𝛾-values with base 10 raised to the power of -5, -4, -3, -2, -1, 0, 1, 2 and 3. The 𝛾-values were not specified in Shahin et al. (2019), thus we defined our own sequence (as discussed in Pedregosa et al., 2011). The optimal hyperparameters found in the grid search were the default (C = 1.0, 𝛾 = ‘scale’ (meaning 𝛾 = 1 / (n_features * variance(X)).

### Appendix A4 – Detailed performance

Table S3: Model trained on Danish triangles and tested on Danish triangles (M1a) - Q1

| actual ↓ predicted→ | **NT** | **ASD** | **Total** |
| --- | --- | --- | --- |
| **NT** | 41 (82 %) | 9 (18 %) | 50 |
| **Autism** | 30 (60 %) | 20 (40 %) | 50 |
| **Total** | 71 | 29 | **100** |

Table S4: Model trained on Danish storytelling and tested on Danish storytelling (M1b) - Q1

| actual ↓ predicted→ | **NT** | **ASD** | **Total** |
| --- | --- | --- | --- |
| **NT** | 10 (100 %) | 0 (0 %) | 10 |
| **Autism** | 2 (25 %) | 6 (75 %) | 8 |
| **Total** | 12 | 6 | **18** |

Table S5: Model trained on US storytelling and tested on US storytelling (M1c) - Q1

| actual ↓ predicted→ | **NT** | **ASD** | **Total** |
| --- | --- | --- | --- |
| **NT** | 23 (82 %) | 5 (18 %) | 28 |
| **Autism** | 16 (50 %) | 16 (50 %) | 32 |
| **Total** | 39 | 21 | **60** |

Table S6: Model trained on Danish triangles and tested on Danish storytelling (M2a) - Q2

| actual ↓ predicted→ | **NT** | **ASD** | **Total** |
| --- | --- | --- | --- |
| **NT** | 41 (87 %) | 6 (13 %) | 47 |
| **Autism** | 20 (40 %) | 30 (60 %) | 50 |
| **Total** | 61 | 36 | **97** |

Table S7: Model trained on Danish storytelling and tested on Danish triangles (M2b) - Q2

| actual ↓ predicted→ | **NT** | **ASD** | **Total** |
| --- | --- | --- | --- |
| **NT** | 164 (68 %) | 77 (32 %) | 241 |
| **Autism** | 100 (37 %) | 172 (63 %) | 272 |
| **Total** | 264 | 249 | **513** |

Table S8: Model trained on Danish storytelling and tested on US storytelling (M3a) - Q3

| actual ↓ predicted→ | **NT** | **ASD** | **Total** |
| --- | --- | --- | --- |
| **NT** | 180 (96 %) | 7 (4 %) | 187 |
| **Autism** | 122 (100 %) | 0 (0 %) | 122 |
| **Total** | 302 | 7 | **309** |

Table S9: Model trained on US storytelling and tested on Danish storytelling (M3b) - Q3

| actual ↓ predicted→ | **NT** | **ASD** | **Total** |
| --- | --- | --- | --- |
| **NT** | 29 (62 %) | 18 (38 %) | 47 |
| **Autism** | 38 (76 %) | 12 (24 %) | 50 |
| **Total** | 67 | 30 | **97** |

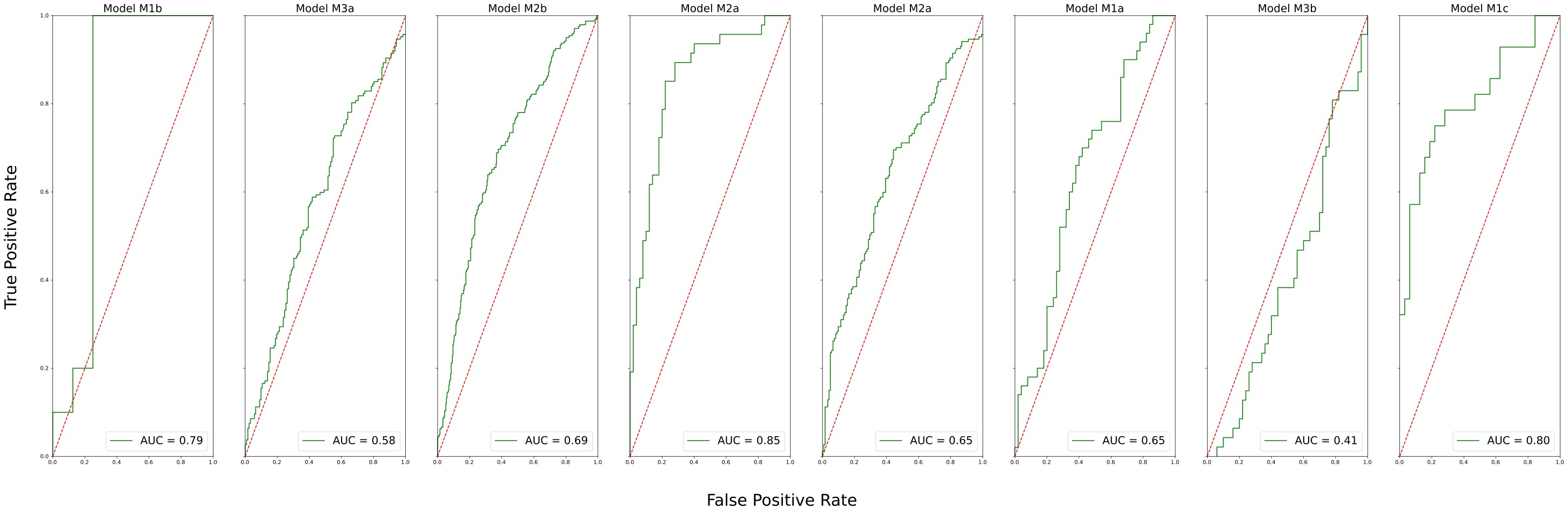

*Figure S1 – Receiving Operator Curve for each model.*

### Appendix A5 – Voice recordings across datasets

*Table S10: number of voice recordings in each dataset. The first three rows are data for the three training sets. The following rows are data for the test sets, which vary in size due to the CV and train/test/hold-out procedures.*

| Dataset | # Total | # NT | # Autism |
| --- | --- | --- | --- |
| --- Train and test on same conditions --- | | | |
| Danish training set - triangle task | 413 | 222 | 191 |
| Danish training set - storytelling task | 79 | 42 | 37 |
| US training set - storytelling task | 249 | 90 | 159 |
| --- Danish test sets --- | | | |
| Tested on Danish storytelling task  Trained on Danish storytelling task | 18 | 10 | 8 |
| Tested on Danish storytelling task  Trained on US storytelling task or Danish triangle task | 97 | 50 | 47 |
| Tested on Danish triangle task  Trained on Danish triangle task | 100 | 50 | 50 |
| Tested on Danish triangle task  Trained on Danish storytelling task | 513 | 272 | 241 |
| --- US test sets --- | | | |
| Tested on US storytelling task  Trained on US storytelling task - | 60 | 32 | 28 |
| Tested on US storytelling task  Trained on Danish storytelling task | 309 | 187 | 122 |
